# Determinants of different facets of beta diversity in Mediterranean marine amphipods

**DOI:** 10.1101/2020.10.30.360677

**Authors:** Bruno Bellisario, Federica Camisa, Chiara Abbattista, Roberta Cimmaruta

## Abstract

Relying on a purely taxonomic view of diversity may ignore the fact that ecological communities can be constituted of species having both distinct evolutionary histories and functional characteristics. Thus, considering how the multiple facets of diversity vary along environmental and geographic gradients may provide insights into the role of historic processes and current environmental conditions in determining the distribution of species, lineages and functions across space. By using distributional, taxonomic-distance and traits information, we explore the role of spatial/environmental gradients and of biogeographic subdivision of Mediterranean Sea on the different facets of beta diversity components in seagrass amphipods. Beta diversity partitioning and correlation analyses showed a nearly equal contribution of the replacement and richness components on total beta diversity for all facets, although the influence of environmental and geographic distance differs among components. While the replacement was mainly related to a pure spatial gradient, both the environmental and geographic distance were correlated with the richness component of beta diversities. Our results are in line with the complex paleobiogeographic history of the Mediterranean Sea, with the replacement component likely to be related to the progressive substitution of species of Atlantic origin with Mediterranean endemics along the west-east geographic gradient, and the richness component to the marked environmental difference between different basins. Moreover, the influence of biogeographic partition on the richness components suggests the role of spatially structured gradients at biogeographic level in determining the net loss/gain of species, lineages and functions, possibly influencing the assembly processes of passive dispersal organisms.

## Introduction

Understanding the patterns and mechanisms of species distribution is a central goal in community ecology, having important outcomes for biodiversity conservation. Although classical approaches focused mainly on the quantification of the difference in the number of species across sites and/or time, recent findings have shown the weakness of considering taxonomic diversity (TD, i.e., species richness) as the unique building block of diversity (Flynn et al. 2011; Cadotte et al. 2012; Craven et al. 2018). Indeed, by using TD as main descriptor of natural communities, one often ignores that the different evolutionary history of species (i.e., their phylogenetic diversity, PD) and the diverse array of functional traits (i.e., their functional diversity, FD) constitute not negligible and complementary facets of diversity (Webb et al. 2002; Petchey and Gaston 2006; Graham and Fine 2008). For instance, distinct communities may be characterized by a high divergence in species composition although showing high levels of functional convergence, signalling the replacement of species performing similar functions across sites (Carvalho et al. 2019). Similarly, functional and phylogenetic diversity may be unrelated as the first usually reflects ecological processes acting at local scales, while the latter is traditionally interpreted as due to the effects of ancient events as biogeographic history (Ramm et al. 2018). Biodiversity investigation taking into account phylogenetic and trait-based analyses can thus help disentangling the mechanisms of community assembly, since differences in their patterns may reveal the signature of processes acting at different spatial and temporal scales.

Beta diversity is a core concept in ecology, aimed at determining the degree of dissimilarity in the assemblage of communities along gradients (Whittaker 1960; Anderson et al. 2011; but see Legendre 2019 and reference therein for a temporal definition of beta diversity). Beta diversity can be decomposed into two additive components of dissimilarity, replacement (or turnover, i.e., the substitution rate of species along the gradient) and richness (i.e., the net difference in species number), each reflecting different processes acting on communities. For instance, the replacement along a given gradient can be interpreted as due to environmental niche filtering (Saladin et al. 2019), while the richness differentiation of assemblages may be due either to the competitive loss of species or dispersal limitation (Si et al. 2016).

Recently, the beta diversity concept has been extended to include other dimensions of diversity besides taxonomical, namely phylogenetic and functional (Graham and Fine 2008; Leprieur et al. 2012; Cardoso et al. 2014). As the total beta diversity and its components may respond differently to ecological and spatial gradients, knowledge of their correlates provides useful information on the factors shaping biodiversity variation (Rocha et al. 2019; Heino et al. 2019). When the effect of pure spatial distance is dominant, differences in community composition are thought to follow neutral models driven by dispersal limitation alone (Hubbel, 2001). Conversely, in case of strong environmental signals, deterministic models based on the ecological responses of species to their surrounding environment (i.e., environmental filtering) should shape beta diversity patterns (Cornwell et al. 2006). However, taxonomic, phylogenetic and functional dissimilarity may result from the joint effect of both geographic and environmental constraints, as environmental variables are intrinsically spatially structured (Legendre and Legendre 2012). Under this circumstance, it can be expected that beta diversity is driven by spatially structured environmental gradients, rather than pure environmental ones (Zhang et al. 2019).

The Mediterranean Sea has physical, chemical and historic features that make it particularly suitable to study the effects of geographic and environmental gradients on beta diversity. It is a concentration basin, where global oceanographic conditions allow for differences in evaporation regime between different portions (higher in the easternmost part), ultimately determining strong environmental gradients mostly related to differences in temperature and salinity (Coll et al. 2010). Moreover, surface circulation coupled with local meteorology and bathymetry, allow for a complex spatial environmental heterogeneity and the development of very steep gradients localized at the boundaries of the main circulation divides (Berline et al. 2014). Such complexity results in well-defined spatial patterns of diversity, generally decreasing from north-western to south-eastern regions, determined by the dispersal ability of species and the selection by local environmental conditions (Coll et al. 2010; Berline et al. 2014).

Within this context, marine benthic amphipods represent a well-suited group of species to test specific hypotheses relating the complementary role of geographic and environmental constraints on the structuring of ecological communities within the Mediterranean Sea. The lack of planktonic larval stages limits the dispersal ability of amphipods, suggesting relatively small distribution ranges, high endemicity and a robust biogeographic pattern (Arfianti and Costello 2020). Understanding the determinants of amphipods distribution over large scales is fundamental, as they represent a key trophic level in benthic ecosystems, being also extremely sensitive to environmental alterations (Pinnegar et al. 2000; Kumagai 2008; Michel et al. 2015). Beta diversity may thus provide useful information on how environmental and/or geographic factors affect the distribution of species, lineages and functions in Mediterranean marine amphipods.

In this paper, we aim to test for the relative contribution of geographic and environmental constraints on the taxonomic, phylogenetic and functional beta diversity. We used distribution data of amphipods associated with *Posidonia oceanica* (L.) Delile, 1813 mined from literature, including information on a selected array of functional traits having important outcomes for the community assembly process: body size, living habit and trophic group. Localities were characterized by using data on the occurrence of *P. oceanica* seagrass habitat and information on specific environmental features (i.e., sea surface temperature and salinity), known to have a direct influence on the distribution of biota in the Mediterranean Sea (Coll et al. 2010). Moreover, as rates of trait evolution and speciation may differ among biogeographic regions (Arnan et al. 2017), we used a previously identified biogeographic partition of the same dataset (Bellisario et al. 2019), to understand the importance of biogeographic barriers with respect to geographic distance and environmental gradients in structuring beta diversity.

### Material and methods Study area

We used data on the presence/absence of 147 amphipod species from *P. oceanica* meadows in 28 sampling localities across the Mediterranean Sea (Fig. 1). The dataset was reconstructed from different sources mined from literature, all selected by comparable data in terms of sampling season, depth and survey methods (see Bellisario et al. 2019 and reference therein). The study area covered a large portion of the Mediterranean Sea, characterized by different geographic, hydrological and geological features, as well as differences in the potential connectivity due to general circulation models (Bianchi 2011; Berline et al. 2014). In a previous work (Bellisario et al. 2019), a network approach based on modularity was applied on the same data set to identify the bioregional partition of local assemblages, identifying four main bioregions corresponding to the main divides of the Mediterranean basin (Fig. 1).

**Figure 1.**
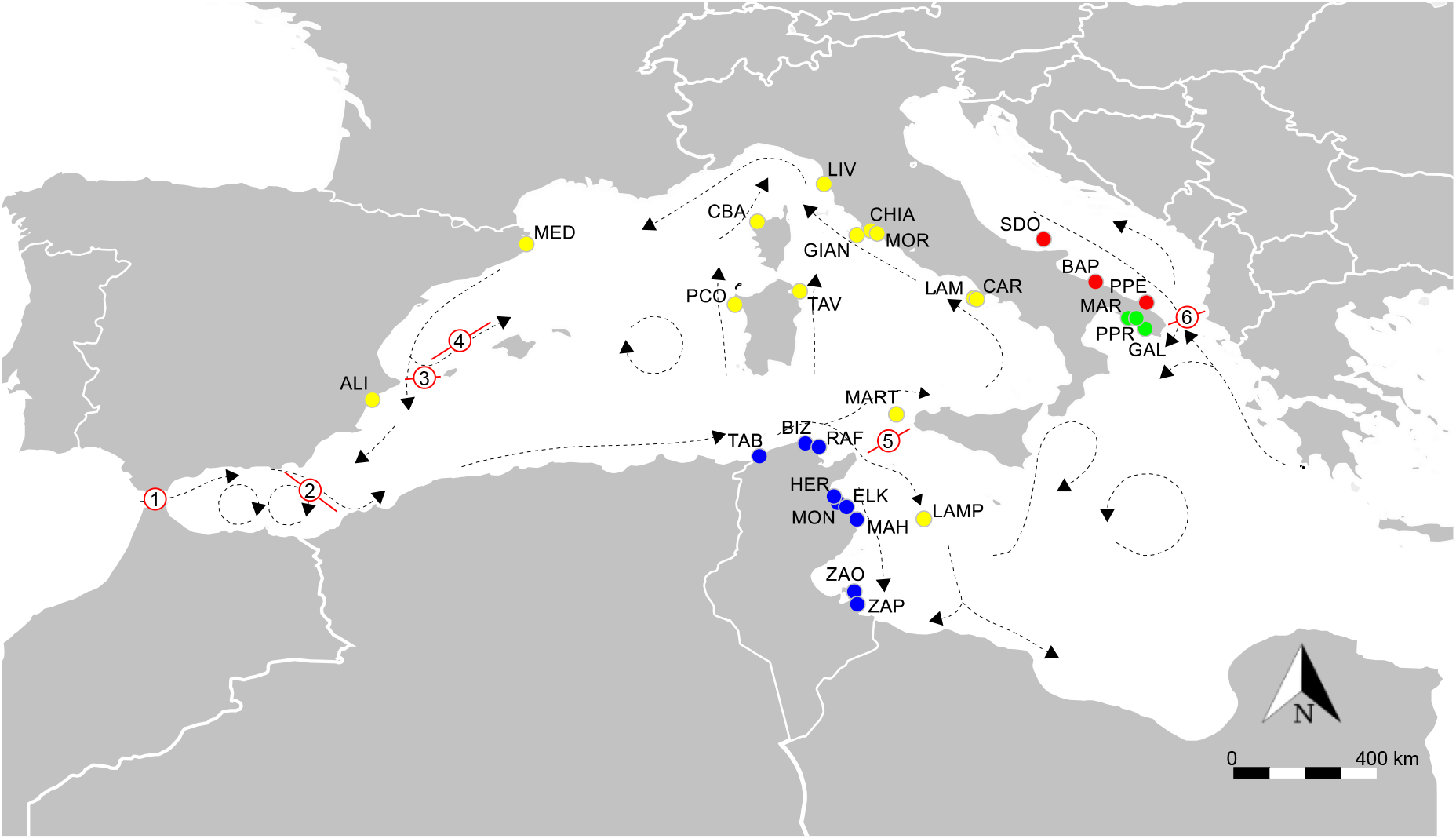
Geographic extension of the study area and sampling localities. Dotted arrows indicate the main circulation patterns and red lines with circles the main barriers: 1, Gibraltar Strait; 2, Almeria-Oran Front; 3, Ibiza Channel; 4, Balearic Front; 5, Sicily Channel; 6, Otranto Strait. Colours correspond to the biogeographic partition: blue, Tunisian (TUN); yellow, Central Western Mediterranean (CWM); red, Adriatic Sea (ADR); green, Ionian Sea (ION). For acronyms, please refer to the Supplementary Materials available in the Data availability statement section.

### Environmental and habitat characterization

We used the Sea Surface Temperature (SST) and Sea Water Salinity (PSU) values as main determinants of the environmental characteristics of sites, since they are both known to co-vary geographically and so influencing the pattern of species diversity throughout the Mediterranean basin (Coll et al. 2010). Data were downloaded from the COPERNICUS Marine Environment Marine Service (CMEMS, https://marine.copernicus.eu), which provides an easy way to access a variety of data from multiple sources (e.g., satellite observations and *in situ* sensors) encompassing different spatial and temporal resolution. SST and PSU were extracted from the Mediterranean Sea Physics Reanalysis, a hydrodynamic model supplied by the Nucleus for European Modelling of the Ocean (NEMO), with a resolution matching the seasonal (Spring/Summer), temporal (1987-2014) and depth coverage (5-20m) of the original papers from which community data were extracted (see Data availability statement section). This model has a variational data assimilation scheme (OceanVAR) for temperature and salinity vertical profiles and satellite sea level anomaly along track data, with an horizontal grid resolution of ∼1 km. Reanalysis has been initialized with a gridded climatology for temperature and salinity computed from *in situ* data sampled before 1987, and released as monthly mean (more information about models and data processing can be found at http://marine.copernicus.eu/services-portfolio/access-to-products/).

To describe the habitat characteristic at sites, we used the probability of *P. oceanica* occurrence (PPOMed), downloaded from the EMODnet Seabed Habitat portal (https://www.emodnet-seabedhabitats.eu) with a spatial resolution of ∼1km and values ranging in the 0-1 interval. PPOMed expresses the probability of seagrass occurrence from the collation and integration of available habitat distribution models at Mediterranean scale, derived by applying a Machine Learning technique (i.e., Random Forest) to train on data from regions where information was available and then used to predict the probability of occurrence of *P. oceanica* (more information can be found in Cameron and Askew 2011).

To derive a meaningfulness measure of the environmental and habitat characteristics, we performed a geostatistical analysis by superimposing a vector of sampling data points to the SST, PSU and PPOMed rasters in a Geographic Information System (QGIS Development Team 2019) and derived the average (± SD) values within a 1km search radius centred on sampling localities.

### Amphipod traits and phylogenetic data

We included information on body size, living habit and trophic groups, which are considered key functional traits determining the competitive ability and community assembly processes of seagrass amphipods (Best et al. 2013; Scipione 2013; Best and Stachowicz 2014; Lürig et al. 2016). Traits information were extracted from available literature on Mediterranean marine amphipods (Ruffo 1982; 1989; 1993; 1998; Scipione 2013) and nomenclature was updated following the World Register of Marine Species (WoRMS, last access 22/02/2021).

Body size was quantified using the maximum body length (measured from the anterior margin of the head to the posterior end of the telson, in mm) in accordance with the fast- seasonal growth cycle of seagrass amphipods (Spring/Summer breeding period, Best and Stachowicz 2014). Since a certain degree of sexual body size dimorphism may exist in amphipods (Longo and Mancinelli 2014), values were taken as the average maximum body length between male and female specimens.

Information on living habits and trophic groups were extracted to the highest possible taxonomic level (i.e., species) and, in case of missing information at species level, we used families as the upper limit for classification. Traits were coded as not available (NA) if no information was available at family level. Following Scipione (2013), each species was thus classified to four living habit categories (epifaunal free-living, epifaunal domicolous, infaunal free-burrowing and infaunal tube-building) and 10 trophic groups (suspension feeders, deposit feeders, carnivores, commensals, herbivores, plant detritus feeders, omnivores, deposit-suspension feeders, deposit feeders-carnivores and herbivores-deposit feeders). Traits distance between species was measured with the Gower distance (Pavoine et al. 2009), which allows for missing trait values and the use of both quantitative (body size) and qualitative (living habit and trophic group) data, by means of the function ‘gowdis’ in the FD package of R (Laliberté and Legendre 2010; R Development Core Team 2018).

Since a true phylogenetic tree was currently unavailable for all amphipod species recorded in our study, we used a taxonomic distance measure based on the average path lengths in the taxonomic tree (Ricotta et al. 2012; Cardoso et al. 2014). Although the amount of approximation provided by surrogate measures for phylogeny poses serious limits when used for answering strict evolutionary questions (Cardoso et al. 2014), their use is found to be valid in large-scale metacommunity studies (Heino and Tolonen 2017; Hill et al. 2019). Taxonomic information were retrieved from WORMS by using the function ‘classification’ in the package ‘taxize’ (Chamberlain and Szocs 2013) of R and the taxonomic distance was calculated by using equal branch lengths between seven taxonomic levels (Species, Genus, Family, Superfamily, Parvorder, Infraorder, Suborder), by using the function ‘taxa2dist’ in the ‘vegan’ package of R (Oksanen et al. 2019).

### Beta diversity estimates

The Jaccard dissimilarity index was chosen as a beta diversity metric reflecting the variation in amphipod assemblage along gradients. Total beta diversity (β_Tot_) was further decomposed into relativised additive fractions of species replacement (β_Repl_) and richness (β_Rich_) components for taxonomic, phylogenetic and functional diversity. Here, we followed the approach proposed by Cardoso et al. (2014), which uses trees as common representation for taxonomic, phylogenetic and functional diversity, according to which part of a global tree is shared by or unique to the compared communities. Traits and phylogenetic distance matrices were converted to trees and then used to quantify the taxonomic (TDβ), phylogenetic (PDβ) and functional (FDβ) beta diversity and its replacement and richness components, by using the package ‘BAT’ of R (Cardoso et al. 2015).

### Statistical analyses

The Mantel and partial Mantel tests (Legendre and Legendre 2012) were used to measure the correlation between the different facets of beta diversity components and between these latter with environmental and geographic variables.

The environmental distance was measured by computing the euclidean distance of PPOMed, SST and PSU between sampling localities. Pairwise geographic distances were calculated with a least-cost paths approach, by using land areas (masked using the European Environment Agency coastline polygon 1:100000) as a barrier in distance calculation. This method allows for a more realistic evaluation of the effective spatial separation than the more commonly used Great Circle or euclidean distances due to the complex geometry of coastlines in the Mediterranean basin (Rattray et al. 2016). We also considered the biogeographic (aka modular) partition already derived from our previous study (Bellisario et al. 2019) to account for the biogeographic-level effects in determining the patterns of dissimilarity. We therefore assigned a numeric code to each site corresponding to the bioregion it belonged (Fig. 1), and then calculated the dissimilarity matrix by means of the euclidean distance. This matrix was used as covariates in the partial Mantel test to account for the biogeographic-level effect on explanatory variables. Data and codes are available from the Data availability statement section.

### Generalized dissimilarity modelling

We used Generalized Dissimilarity Modelling (GDM, Ferrier et al. 2007) to model the relationship of TDβ, PDβ and FDβ components with environmental variables. GDM is a statistical regression technique able to accommodate for nonlinearity in eco-geographical datasets, where the compositional dissimilarity provided by beta diversity estimates is modelled as a nonlinear function of the environmental distance between pairs (Ferrier et al. 2007). Moreover, GDM is known to be robust to multicollinearity among predictor variables (e.g., Glassman et al. 2017), facilitating the understanding of the variation in beta diversity along actual environmental gradients.

Here, we discarded the geographic distance provided by latitudinal and longitudinal coordinates as poorly representative of the real geographic distance in our system. Thus, we fitted GDMs using three predictor variables, PPOMed, SST and PSU and plot the I-splines (i.e., monotone cubic spline functions) to assess the impact of predictor variables on the total, replacement and richness components of taxonomic, phylogenetic and functional dissimilarity matrices. The slope of the I-splines curves indicates the rate of dissimilarity while the maximum height represents the total amount of dissimilarity associated with the variable, holding all other variables constant. Variable’s importance was estimated from the sum of each I-spline coefficient. Here, we used the default setting of three I-splines for each predictor, using the package ‘gdm’ of R (Manion et al. 2018).

## Results

### Beta diversity patterns

Assemblages of Mediterranean seagrass amphipods showed relatively high values of total beta diversity for TDβ, slightly lower for PDβ and exhibited the lowest value for FDβ (TDβ_Tot_ = 0.799 ± 0.014; PDβ_Tot_ = 0.624 ± 0.011; FDβ_Tot_ = 0.509 ± 0.009). The replacement and richness components accounted for by a nearly equal contribution for all beta diversity facets (TDβ_Repl_ = 0.418 ± 0.007, TDβ_Rich_ = 0.381 ± 0.006; PDβ_Repl_ = 0.298 ± 0.005, PDβ_Rich_ =0.326 ± 0.006; FDβ_Repl_ = 0.232 ± 0.004, FDβ_Rich_ = 0.277 ± 0.005). As expected, we found significant and positive correlations between the different facets and components of beta diversity (R^2^ > 0.8 and p < 0.001 for all cases). Correlations between the components of PDβ and FDβ still remained significant, although lower, after controlling for TDβ (R^2^ > 0.45 and p < 0.001 for all cases) (Fig. SM1 in Supplementary Materials).

### Correlates of beta diversity

Mantel and partial Mantel tests did not show substantial differences between the total beta diversity of different facets, which resulted to be always positively correlated with both environmental and geographic distances, even after controlling for the biogeographic partition (Table 1). Conversely, the replacement and richness components showed different patterns with respect to the environmental and geographic gradients.

**Table 1.**
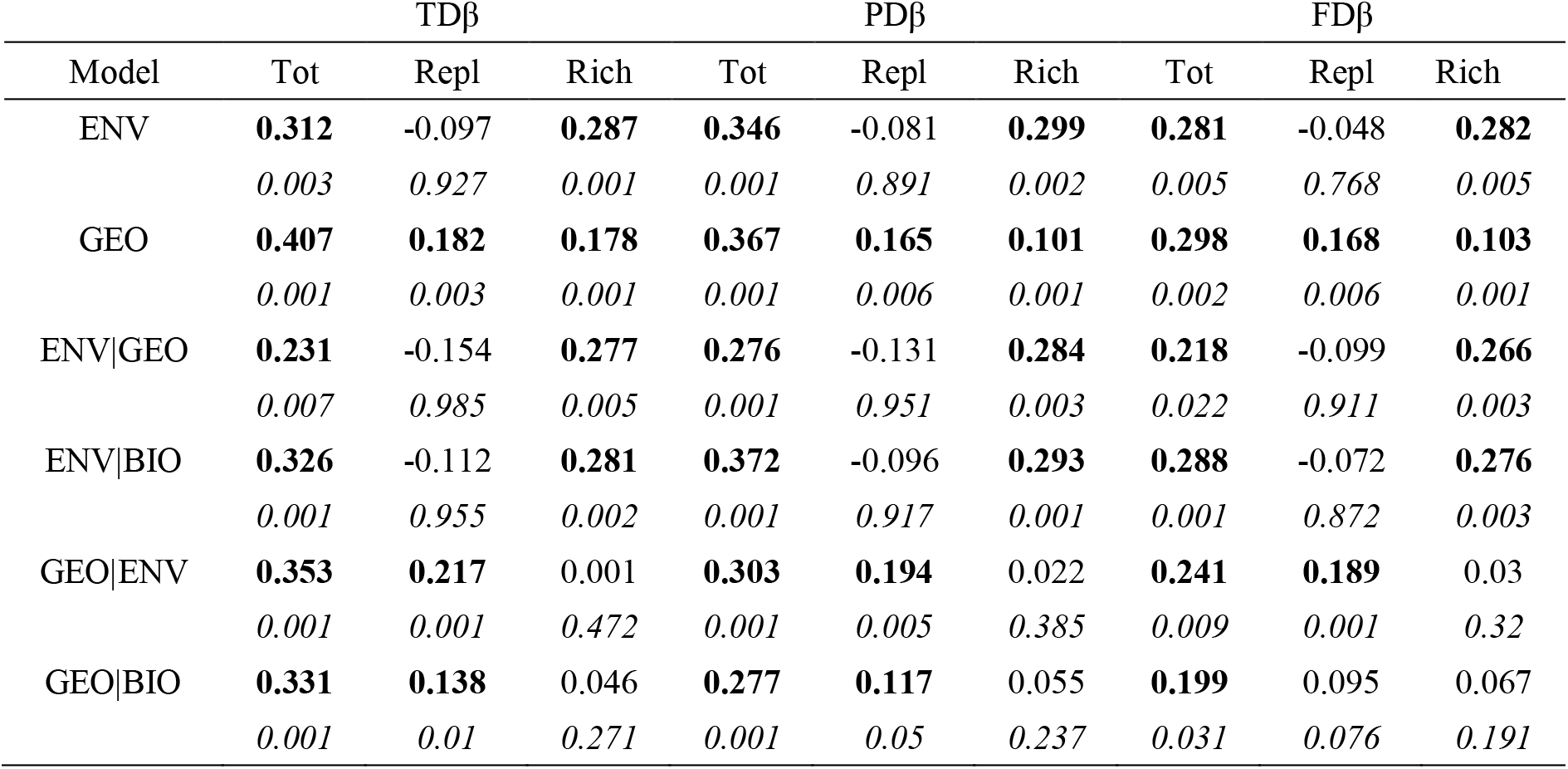
Results of the (partial) Mantel tests between taxonomic (TDβ), phylogenetic (PDβ) and functional (FDβ) beta diversity components of Mediterranean seagrass amphipods and environmental (ENV) and geographic (GEO) distances, while accounting for the biogeographic-level (BIO) effect (see Fig. 1). Bold is for significant values, with p values shown in italic.

| Model | TDβ | | | PDβ | | | FDβ | | |
| --- | --- | --- | --- | --- | --- | --- | --- | --- | --- |
|  | Tot | Repl | Rich | Tot | Repl | Rich | Tot | Repl | Rich |
| ENV | **0.312**<br>*0.003* | -0.097<br>*0.927* | **0.287**<br>*0.001* | **0.346**<br>*0.001* | -0.081<br>*0.891* | **0.299**<br>*0.002* | **0.281**<br>*0.005* | -0.048<br>*0.768* | **0.282**<br>*0.005* |
| GEO | **0.407**<br>*0.001* | **0.182**<br>*0.003* | **0.178**<br>*0.001* | **0.367**<br>*0.001* | **0.165**<br>*0.006* | **0.101**<br>*0.001* | **0.298**<br>*0.002* | **0.168**<br>*0.006* | **0.103**<br>*0.001* |
| ENV\|GEO | **0.231**<br>*0.007* | -0.154<br>*0.985* | **0.277**<br>*0.005* | **0.276**<br>*0.001* | -0.131<br>*0.951* | **0.284**<br>*0.003* | **0.218**<br>*0.022* | -0.099<br>*0.911* | **0.266**<br>*0.003* |
| ENV\|BIO | **0.326**<br>*0.001* | -0.112<br>*0.955* | **0.281**<br>*0.002* | **0.372**<br>*0.001* | -0.096<br>*0.917* | **0.293**<br>*0.001* | **0.288**<br>*0.001* | -0.072<br>*0.872* | **0.276**<br>*0.003* |
| GEO\|ENV | **0.353**<br>*0.001* | **0.217**<br>*0.001* | 0.001<br>*0.472* | **0.303**<br>*0.001* | **0.194**<br>*0.005* | 0.022<br>*0.385* | **0.241**<br>*0.009* | **0.189**<br>*0.001* | 0.03<br>*0.32* |
| GEO\|BIO | **0.331**<br>*0.001* | **0.138**<br>*0.01* | 0.046<br>*0.271* | **0.277**<br>*0.001* | **0.117**<br>*0.05* | 0.055<br>*0.237* | **0.199**<br>*0.031* | 0.095<br>*0.076* | 0.067<br>*0.191* |

The replacement component of all beta diversity facets was correlated with geographic distance and such correlations were robust enough even after partialing out, with the only exception of FDβ_Repl_, which showed no correlation with the geographic distance after controlling for the biogeographic partition (Table 1). No correlation was detected with environmental distance, signalling that the replacement was not influenced by the environment (Table 1). The richness component of beta diversities was correlated with both the environmental and geographic distance, although this latter lacked a correlation when considering the effect of either the environment or biogeography (Table 1). Conversely, the environmental distance still remained significantly correlated when accounting for the geographic and biogeographic distance (Table 1).

GDMs showed substantial differences between the components of each beta diversity facet when considering environmental predictors, being relatively higher for the total beta diversity and showing a negligible contribution on the replacement component with respect to the richness one (Table 2). TDβ_Tot_ was mostly impacted by salinity (Table 2), which showed a linear relationship and then exhibited a sudden increase approximately around 38 PSU, after which the species compositional variation increased rapidly (Fig. 2c). The same trend was observed for PDβ_Tot_ and FDβ_Tot_, although the contribution of PSU was comparable to that of other predictors (Table 2), and the shape of relationship showed the same pattern observed for TDβ_Tot_, although less marked (Fig. 2c). About the replacement component, none of beta diversity facets was impacted by PPOMed (Table 2 and Fig. 2d). SST had a relatively high impact on FDβ_Repl_, while PSU impacted most TDβ_Repl_, and both had a relatively similar effect on PDβ_Repl_ (Table 2). For both FDβ_Repl_ and PDβ_Repl_, a sharp increase was observed within a narrow range of temperature between 17-18 °C, followed by a slow increase for higher temperatures, while the impact on TDβ_Repl_ was mostly linear with a moderate slope (Fig. 2e). Both taxonomic and phylogenetic replacement showed a curvilinear response to PSU, while FDβ_Repl_ exhibited a linear relationship (Fig. 2f). Considering the richness component, PPOMed provided the highest impact on all the facets, all showing a curvilinear relationship followed by a plateau (Fig. 2h). SST had a minor impact on all facets (Fig. 2i), while PSU showed a more pronounced impact characterized by no effect until the threshold of about 38 PSU, beyond which the taxonomic, phylogenetic and functional richness component of beta diversity showed a sudden increase (Fig. 2l).

**Table 2.** GDM model summary for each facets and component of beta diversity with respect to environmental variables.

| | TDβ<sub>Tot</sub> | TDβ<sub>Repl</sub> | TDβ<sub>Rich</sub> | PDβ<sub>Tot</sub> | PDβ<sub>Repl</sub> | PDβ<sub>Rich</sub> | FDβ<sub>Tot</sub> | FDβ<sub>Repl</sub> | FDβ<sub>Rich</sub> |
| --- | --- | --- | --- | --- | --- | --- | --- | --- | --- |
| **Gradient** |  |  |  |  |  |  |  |  |  |
| PPOMed | 0.551 | 0 | 0.444 | 0.373 | 0 | 0.355 | 0.183 | 0 | 0.232 |
| SST | 0.646 | 0.087 | 0.099 | 0.31 | 0.102 | 0.0877 | 0.404 | 0.177 | 0.078 |
| PSU | 1.408 | 0.282 | 0.294 | 0.468 | 0.12 | 0.243 | 0.455 | 0.0988 | 0.233 |
| **GDM summary** |  |  |  |  |  |  |  |  |  |
| Model deviance (%) | 28.727 | 79.244 | 77.411 | 19.586 | 56.971 | 61.931 | 23.109 | 53.305 | 50.268 |
| NULL deviance | 44.529 | 79.759 | 91.811 | 28.391 | 58.374 | 74.331 | 34.339 | 55.672 | 61.353 |
| Deviance explained (%) | 35.481 | 4.407 | 15.685 | 31.011 | 2.403 | 16.682 | 32.701 | 4.252 | 18.066 |

**Figure 2.**
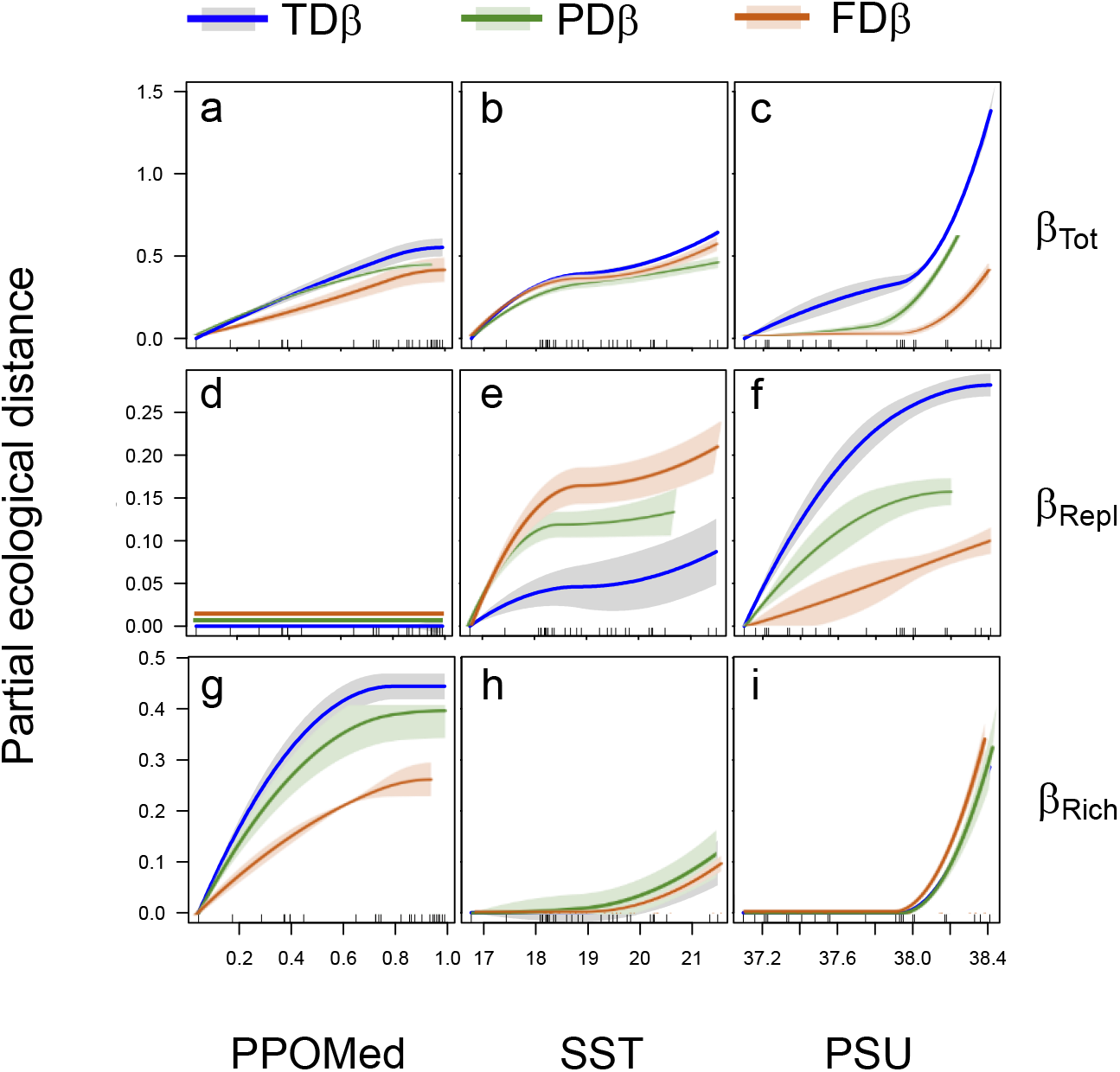
Plots of I-splines of the environmental predictors and confidence intervals from bootstrapping (shaded colours) for the beta diversity components of different facets.

## Discussion

In this study, we focused on the beta diversity of Mediterranean seagrass amphipods to disentangle the role of environmental and geographic gradients on taxonomic, phylogenetic and functional facets of diversity. FDβ showed the lowest value of total beta diversity with respect to both TDβ and PDβ, but a nearly equal contribution of both the replacement and richness components was found for all the three facets considered. However, environmental and geographic gradients influenced differently the components of each facet. Geographic distance was the only determinant of the replacement component, while environmental distance and, to a lesser extent, spatial distance, mainly influenced the richness component. Moreover, the not negligible role of biogeographic partition found for the richness components suggest the role of spatially structured gradients in determining the net loss/gain of species, lineages and functions in Mediterranean seagrass assemblages.

Taxonomic and phylogenetic beta diversity are usually associated with events occurring on wide scales and over long time, typically within a biogeographic frame. The same vision interprets functional beta diversity as mainly, although not exclusively, due to local ecological processes such as environmental filtering (Ramm et al. 2018; Pavoine and Bonsall 2011). Accordingly, different factors might have driven the dynamics of current distribution of species and lineages in seagrass amphipods to produce the observed patterns, mainly related to both the complex paleogeographic history and the marked biogeographic structure of the Mediterranean Sea (Bianchi et al. 2011). About the Mediterranean history, it is well known the role of geological events occurred during the Tertiary and the climatic fluctuations during the Quaternary (especially the most recent cycles of Plio-Pleistocene glaciations; Coll et al. 2010) in determining the distribution of biota in the basin (Bianchi et al. 2011). Moreover, the turbulent geological history of the basin allowed for the creation of a great variety of climatic and hydrologic conditions in fairly isolated sub-basins, subdividing the Mediterranean Sea in many biogeographic sectors characterized by different environmental and habitat features (Bianchi et al. 2011). This allowed species of different biogeographic origin to enter and settle within the basin, contributing to the high level of α-diversity and endemism rate designating the Mediterranean Sea as one of the world’s biodiversity hotspots (Lejeusne et al. 2010; Bianchi et al. 2011).

This scenario is compatible with the high level of β-diversity found in seagrass amphipods in our study and with the balanced contribution of its components. Indeed, replacement could be related to the progressive substitution of species of Atlantic origin with Mediterranean endemics along the west-east axis and, to a lesser extent, along the north-south axis (Bianchi et al. 2011) so following a geographic gradient, while the richness difference could be due to the extremely low environmental affinities of the eastern sectors (i.e., Adriatic and Ionian Sea) with the central-western part of the basin. This hypothesis is in line with the reduced occurrence of Mediterranean endemics and number of shared species with other biogeographic sectors observed in the eastern areas (Bianchi et al. 2011; Bellisario et al. 2019). Indeed, it has been shown that 95% of the known species of Mediterranean amphipods can be recovered in the Central basin, while only 53% inhabit the Adriatic Sea (Bellan-Santini and Ruffo 2003). Thus, the balanced contribution of the replacement and richness on the overall dissimilarity would stem from the combined action of: i) repeated isolation and contraction of biotas and associated speciation events, which might have contributed to the overall diversity of taxa and the divergence of lineages and, ii) variation in environmental features, able to filter which species can survive in particularly selective environments as, for instance, the Adriatic Sea.

This scenario is also suitable for the functional diversity, that however showed a higher influence of the biogeographic subdivision with respect to the geographic cline. Specific features of the Mediterranean circulation and bioregional subdivision can help explain this finding, together with the low dispersal potential of seagrass amphipods. Indeed, a recent eco-regionalization based on the potential connectivity assessed from ensemble Lagrangian simulations provided an in-depth subdivision of the basin in several different regions whose hydrodynamical boundaries can help explain the spatial distribution of passively transported organisms (Berline et al. 2014). Biogeographic boundaries largely match the major discontinuities in variables describing the environment and geographic clines in temperature and salinity characterizing the Mediterranean basin show sharp changes at the main divides, resulting in geographically adjoining but ecologically dissimilar regions (Coll et al. 2011, Berline et al. 2014).

Under this scenario, environmental discontinuities among bioregions might be involved in determining the spatial distribution of functional traits, by sorting species according to their environmental and habitat requirements. For instance, significant differences in the body size distribution across bioregions (but not of living habits and trophic groups) have shown how the Adriatic bioregion is composed of species significantly larger than other bioregions (Fig. SM2, Supplementary Materials). Interestingly, body size is considered a key trait related to the community assembly processes of seagrass amphipods, being involved in a series of complex relationships occurring between the abiotic features of surrounding environment and direct and indirect biotic interactions, mainly due to competition and predation avoidance. Body size, temperature and salinity are dominant factors affecting the metabolic rate in amphipods (Poulin and Hamilton 1995; Maranhão and Marques 2003), which may however vary significantly in response to different ecological condition as, for instance, predation pressure (Glazier et al. 2020). Moreover, historical and biogeographic processes, alongside current environmental conditions, may have played a role in determining the distribution and complexity of seagrass habitat (*sensu* Hacker and Steneck 1990), thereby allowing for the functional differentiation of assemblages at bioregional scale as response to an evolved consequence of consistently high predation risk, size-dependent habitat selection and food availability (Kovalenko et al. 2011; Lürig et al. 2016).

GDM analysis supports these findings, highlighting the impact of salinity and the presence of a threshold at 38 PSU, beyond which beta diversity metrics showed an exponential increase (Fig. 2). This value corresponds to the southern Adriatic/Ionian Sea surface salinity, and is abruptly reached by crossing the Sicily Channel, a main divide of the Mediterranean Sea. In the case of Adriatic Sea, environmental features are coupled with the presence of the strong barrier represented by the Strait of Otranto, and previous studies have shown that this area is inhabited exclusively by widely distributed amphipod species, mainly cosmopolitan, as a result of both extreme environmental conditions and geographic isolation (Bellisario et al., 2019). Moreover, such threshold sets also the tolerance limit for the activation of osmoregulatory processes that may potentially interfere with the leaf growth, survival and photosynthetic rates of *P. oceanica*, thereby affecting meadow structure and limiting the distribution of the seagrass in the easternmost sectors of the Mediterranean (Sandoval-Gil et al. 2012).

In conclusion, our findings show how taxonomic, phylogenetic and functional beta diversity in Mediterranean seagrass amphipods stem from an equal contribution of both the replacement and richness components, which however can be driven by distinct processes with respect to the beta diversity facets analysed. While geographic distance alone is the main constraint determining the replacement, spatially structured gradients at biogeographic scale mainly determine the net loss/gain of species, lineages and functions. Overall, our results corroborate the hypothesis that, although the Mediterranean Sea is largely characterized by wide-basin gradients, biogeographic boundaries may have a strong influence in accounting for biodiversity distribution (Bianchi et al. 2011), by creating marked discontinuities in environmental and spatial gradients, possibly influencing dispersal- and niche-based processes in the community assembly of passive dispersal organisms.

## Declarations

### Availability of data and material

Data and codes are available from Supplementary Materials.

### Authors’ contributions

BB and RC conceived and designed the paper, wrote the manuscript and approved the final draft; BB analysed the data, prepared figures and tables; FC and CA collected field materials. All authors gave final approval for publication.

